# Metabolites associated with early cognitive changes implicated in Alzheimer’s disease

**DOI:** 10.1101/436667

**Authors:** Burcu F. Darst, Erin M. Jonaitis, Rebecca L. Koscik, Lindsay R. Clark, Qiongshi Lu, Kirk J. Hogan, Sterling C. Johnson, Corinne D. Engelman

**Affiliations:** University of Wisconsin, Madison, 333 East Campus Mall, Madison, WI, 53706, USA; Department of Population Health Sciences, University of Wisconsin School of Medicine and Public Health, 610 Walnut Street, Madison, WI, 53726, USA; Wisconsin Alzheimer’s Institute, University of Wisconsin School of Medicine and Public Health, 610 Walnut Street, Madison, WI, 53719, USA; Geriatric Research Education and Clinical Center, Wm. S. Middleton Memorial VA Hospital, 2500 Overlook Terrace, Madison, WI, 53705, USA; Alzheimer’s Disease Research Center, University of Wisconsin School of Medicine and Public Health, 600 Highland Avenue, Madison, WI, 53792, USA; Department of Biostatistics & Medical Informatics, University of Wisconsin, Madison, 600 Highland Avenue, Madison, WI, 53792, USA; Department of Anesthesiology, University of Wisconsin School of Medicine and Public Health, 600 Highland Avenue, Madison, Madison, WI, 53792, USA

**Keywords:** Metabolomics, cognition, Alzheimer’s disease, fatty acids, amino acids, longitudinal analysis

## Abstract

We investigated the metabolomics of early cognitive changes related to Alzheimer’s disease (AD) in order to better understand mechanisms that could contribute to early stages and progression of this disease. This investigation used longitudinal plasma samples from the Wisconsin Registry for Alzheimer’s Prevention (WRAP), a cohort of participants who were dementia free at enrollment and enriched with a parental history of AD. Metabolomic profiles were quantified for 2,338 fasting plasma samples among 1,206 participants, each with up to three study visits. Of 1,097 metabolites tested, levels of seven were associated with executive function trajectories, including an amino acid and three fatty acids, but none were associated with delayed recall trajectories. Our time-varying metabolomic results suggest potential mechanisms that could contribute to the earliest signs of cognitive decline. In particular, fatty acids may be associated with cognition in a manner that is more complex than previously suspected.

## 1. Introduction

Recent technological advances have made metabolomic studies increasingly favorable among Alzheimer’s disease (AD) researchers (Enche Ady et al., 2017; Gonzalez-Dominguez et al., 2017; Trushina and Mielke, 2014); however, most of these studies have been limited to cross-sectional approaches comparing patients with either AD or mild cognitive impairment (MCI) to controls. In these early stages of AD metabolomics research, few metabolites have been found to be associated with AD in more than one study (Enche Ady et al., 2017). Because neuropathological changes that lead to the development of AD occur decades before its clinical presentation (Jack et al., 2010; Serrano-Pozo et al., 2011), longitudinal investigations preceding its diagnosis could add to our current knowledge. In particular, understanding how biomarkers correlate with subtle changes in cognition prior to AD diagnosis could help identify causal mechanisms contributing to its onset.

Executive function and memory deficits occur in the very early stages of AD, prior to deficits of language and visuospatial functions (Albert, 1996; Baudic et al., 2006; Lafleche and Albert, 1995), and are associated with AD pathology and subsequent global cognitive decline (Clark et al., 2016; Clark et al., 2012). Metabolite levels associated with these early changes in cognition could be indicative of underlying biological mechanisms and pathways contributing to the pathology of AD and could ultimately inform stronger predictive models for this disease.

Using longitudinal plasma samples from the Wisconsin Registry for Alzheimer’s Prevention (WRAP), we investigated whether time-varying metabolite levels predicted age-related cognitive changes (*i.e.*, trajectories) for executive function and memory, specifically, delayed recall. Results from each of these association analyses were further explored using Mendelian randomization (MR) and in a metabolite pathway analysis.

## 2. Methods

### 2.1. Participants

Study participants were from WRAP, a longitudinal study of initially dementia free middle-aged adults that allows for the enrollment of siblings and is enriched for a parental history of AD. Further details of the study design and methods used have been previously described (Johnson et al., 2018; Sager et al., 2005). Analyses did not include the baseline WRAP visit due to subsequent protocol changes regarding sample collection procedures and tests included in the neuropsychological battery. Study visits included in the current analyses occurred every two years, with plasma samples and cognitive measures collected concurrently within the same study visit. This study was conducted with the approval of the University of Wisconsin Institutional Review Board, and all participants provided signed informed consent before participation.

### 2.2. Biological Samples

#### 2.2.1. Plasma collection and sample handling

Fasting blood samples for this study were drawn the morning of each study visit, which was also the day cognitive testing was completed. Blood was collected in 10 mL ethylenediaminetetraacetic acid (EDTA) vacutainer tubes. They were immediately placed on ice, and then centrifuged at 3000 revolutions per minute for 15 minutes at room temperature. Plasma was pipetted off within one hour of collection. Plasma samples were aliquoted into 1.0 mL polypropylene cryovials and placed in −80°C freezers within 30 minutes of separation. Samples were never thawed before being shipped overnight on dry ice to Metabolon, Inc. (Durham, NC), where they were again stored in −80°C freezers and thawed once before testing.

#### 2.2.2. Metabolomic profiling and quality control

An untargeted plasma metabolomics analysis was performed by Metabolon, Inc. using Ultrahigh Performance Liquid Chromatography-Tandom Mass Spectrometry (UPLC-MS/MS). Quantification was performed as previously described (Evans et al., 2014); details are outlined in the Supplemental Note. Metabolites within nine super pathways were identified: amino acids, carbohydrates, cofactors and vitamins, energy, lipids, nucleotides, partially characterized molecules, peptides, and xenobiotics.

Up to three longitudinal plasma samples were available for each participant. Metabolites with an interquartile range of zero (*i.e.*, those with very low or no variability due to individuals having almost identical levels of the given metabolite) were excluded from analyses (n=178 metabolites). After removing these metabolites, samples were missing a median of 11.7% metabolites, while metabolites were missing in a median of 1.2% of samples. Missing metabolite values were imputed to the lowest level of detection for each metabolite. Metabolite values were median-scaled and log-transformed to normalize metabolite distributions (van den Berg et al., 2006). If a participant reported that they did not fast or withhold medications and caffeine for at least eight hours, the sample was excluded from analyses (n=159 samples). A total of 1,097 metabolites among 2,338 samples remained for analyses.

#### 2.2.3. DNA collection and genomics quality control

DNA was extracted from whole blood samples using the PUREGENE^®^ DNA Isolation Kit (Gentra Systems, Inc., Minneapolis, MN). DNA concentrations were quantified using the Invitrogen™ Quant-iT™ PicoGreen™ dsDNA Assay Kit (Thermo Fisher Scientific, Inc., Hampton, NH) analyzed on the Synergy 2 Multi-Detection Microplate Reader (Biotek Instruments, Inc., Winooski, VT). Samples were diluted to 50 ng/ul following quantification.

A total of 1,340 samples were genotyped using the Illumina Multi-Ethnic Genotyping Array at the University of Wisconsin Biotechnology Center (Figure S1). Thirty-six blinded duplicate samples were used to calculate a concordance rate of 99.99%, and discordant genotypes were set to missing. Sixteen samples missing >5% of variants were excluded, while 35,105 variants missing in >5% of individuals were excluded. No samples were removed due to outlying heterozygosity. Six samples were excluded due to inconsistencies between self-reported and genetic sex.

Due to sibling relationships in the WRAP cohort, genetic ancestry was assessed using Principal Components Analysis in Related Samples (PC-AiR), a method that makes robust inferences about population structure in the presence of relatedness (Conomos et al., 2015). This approach included several iterative steps and was based on 63,503 linkage disequilibrium (LD) pruned (r^2^<0.10) and common (MAF>0.05) variants, using the 1000 Genomes data as reference populations (Genomes Project et al., 2015). First, kinship coefficients (KCs) were calculated between all pairs of individuals using genomic data with the Kinship-based Inference for Gwas (KING)-robust method (Manichaikul et al., 2010). PC-AiR was used to perform principal components analysis (PCA) on the reference populations along with a subset of unrelated individuals identified by the KCs. Resulting principal components (PCs) were used to project PC values onto the remaining related individuals. All PCs were then used to recalculate the KCs taking ancestry into account using the PC-Relate method, which estimates KCs robust to population structure (Conomos et al., 2016). PCA was performed again using the updated KCs, and KCs were also estimated again using updated PCs. The resulting PCs identified 1,198 WRAP participants whose genetic ancestry was primarily of European descent. This procedure was repeated within this subset of participants (excluding 1000 Genomes individuals) to obtain PC estimates used to adjust for population stratification in subsequent genomic analyses. Among European descendants, 160 variants were not in Hardy-Weinberg equilibrium (HWE) and 327,064 were monomorphic and thus, removed.

A total of 1,294,660 bi-allelic autosomal variants among 1,198 European descendants remained for imputation, which was performed with the Michigan Imputation Server v1.0.3 (Das et al., 2016), using the Haplotype Reference Consortium (HRC) v. r1.1 2016 (McCarthy et al., 2016) as the reference panel and Eagle2 v2.3 (Loh et al., 2016) for phasing. Prior to imputation, the HRC Imputation Checking Tool (Rayner et al., 2016) was used to identify variants that did not match those in HRC, were palindromic, differed in MAF>0.20, or that had non-matching alleles when compared to the same variant in HRC, leaving 898,220 for imputation. A total of 39,131,578 variants were imputed. Variants with a quality score R^2^<0.80, MAF<0.001, or that were out of HWE were excluded, leaving 10,400,394 imputed variants. These were combined with the genotyped variants, leading to 10,499,994 imputed and genotyped variants for analyses. Data cleaning and file preparation were completed using PLINK v1.9 (Chang et al., 2015) and VCFtools v0.1.14 (Danecek et al., 2011). Coordinates are based on GRCh37 assembly hg19.

### 2.3. Cognitive phenotypes

Composite scores were calculated for executive function and delayed recall based on a previous analysis (Clark et al., 2016). Each composite score was calculated from three neuropsychological tests, which were each converted to z-scores using baseline means and standard deviations. The executive function composite score included the Trails Making Test Part B (TMTB) (Reitan and Wolfson, 1985) total time to completion, Stroop Neuropsychological Screen Test (Trenerry et al., 1989) color-word interference total items completed in 120 second, and Wechsler Abbreviated Intelligence Scale-Revised (WAIS-R) digit symbol coding subtest total items completed in 90 seconds (Wechsler, 1981). The delayed recall composite score included the Rey Auditory Verbal Learning Test (RAVLT) (Schmidt, 1996) long-delay free recall, Wechsler Memory Scale-Revised Logical Memory (WMS-R LM) (Wechsler, 1987) delayed recall, and Brief Visuospatial Memory Test (BVMT-R) (Benedict, 1997) delayed recall. The TMTB was multiplied by negative one prior to being converted to z-scores, so that higher z-scores indicated better performance. These z-scores were then averaged to derive executive function and delayed recall composite scores at each visit for each individual.

### 2.4. Statistical analyses

#### 2.4.1. Metabolome-wide association studies

All associations were tested using linear mixed effects regression models implemented in the SAS MIXED procedure. To assess whether metabolite levels were associated with age-related cognitive trajectories, an interaction term between metabolite level and age was used to predict cognitive composite scores (*i.e.*, executive function and delayed recall). Models included fixed effects for centered age, sex, self-reported race, cholesterol-lowering medication use (the most commonly used class of medications in our sample), sample storage time, education, a genetic risk score for *APOE* (Darst et al., 2017), and practice effects (using visit number). Random intercepts were included for within-subject correlations due to repeated measures and within-family correlations due to the enrollment of siblings. The two sets of *P*-values resulting from testing executive function and delayed recall trajectories were separately corrected for multiple testing using the Benjamini-Hochberg (Benjamini and Hochberg, 1995) adjustment with an alpha of 0.05.

#### 2.4.2. Mendelian randomization

MR (Smith and Ebrahim, 2003) was used to assess whether levels of any individual metabolite identified in our association analyses (*i.e.*, metabolites associated with either executive function or delayed recall trajectories) could causally influence cognition. Metabolic quantitative trait loci (mQTL) were identified as genomic variants influencing metabolite levels with a *P*<0.001 using genome-wide association study (GWAS) summary statistics provided by the authors of a recent publication by Long et al., 2017 (Long et al., 2017). A polygenic score (PS) was created for each metabolite identified in our association analyses that also had GWAS summary statistics available. PSs were defined as the sum of an individual’s metabolite-increasing alleles weighted by the effect sizes from GWAS summary statistics. PSs were created using the additive allelic scoring function in PLINK 1.9 (Chang et al., 2015) after LD pruning variants within each PS (R^2^>0.50). To be consistent with our discovery models, interactions between each PS and age were tested for association with cognition using linear mixed effects regression models. Models included fixed effects for age, sex, education, practice effects, and the first four PCs to account for population stratification. They also included random intercepts for repeated measures and sibling relationships.

#### 2.4.3. Metabolite pathway analysis

Results from association analyses were further investigated using a metabolite pathway analysis. Metabolites included in this analysis were those associated with either executive function or delayed recall trajectories with an unadjusted *P*<0.05 and that had a Kyoto Encyclopedia of Genes and Genomics (KEGG) compound ID (Kanehisa and Goto, 2000). Metabolites on our panel with KEGG compound IDs were used as the reference panel for this analysis. The pathway analysis was conducted using MetaboAnalyst 4.0 and included both an overrepresentation analysis, which was assessed using a hypergeometric test, and a pathway topology analysis, which was assessed using relative-betweenness centrality (Xia and Wishart, 2016). The overrepresentation analysis tests whether a user-defined list of metabolites represents a particular pathway of metabolites more than expected by chance. The pathway topology analysis considers the structure of a pathway by assessing how connected metabolites are within a pathway. If a pathway contains metabolites that connect dense clusters of other metabolites, the pathway would have a high impact score, as changes to its metabolites would likely have a strong impact on other metabolites within the pathway.

### 2.5. Data Availability

Data are available on request made to the WRAP Science Executive Committee. Details and application instructions can be found using the following website: http://www.wai.wisc.edu/research/wrapdatarequests.html.

## 3. Results

### 3.1. Participants

A total of 1,206 WRAP participants with 2,338 longitudinal plasma samples were available for analyses. At baseline for the current study, 69.2% of participants were female, 93.7% were Caucasian, and participants were 60.9 years old with a bachelor’s degree, on average (Table 1). Participants each had 1,097 plasma metabolites available for analyses, 347 (31.6%) of which were of unknown chemical structure. Properties of each metabolite, such as biochemical name, super pathway, and sub pathway are described in Table S1.

**Table 1.**
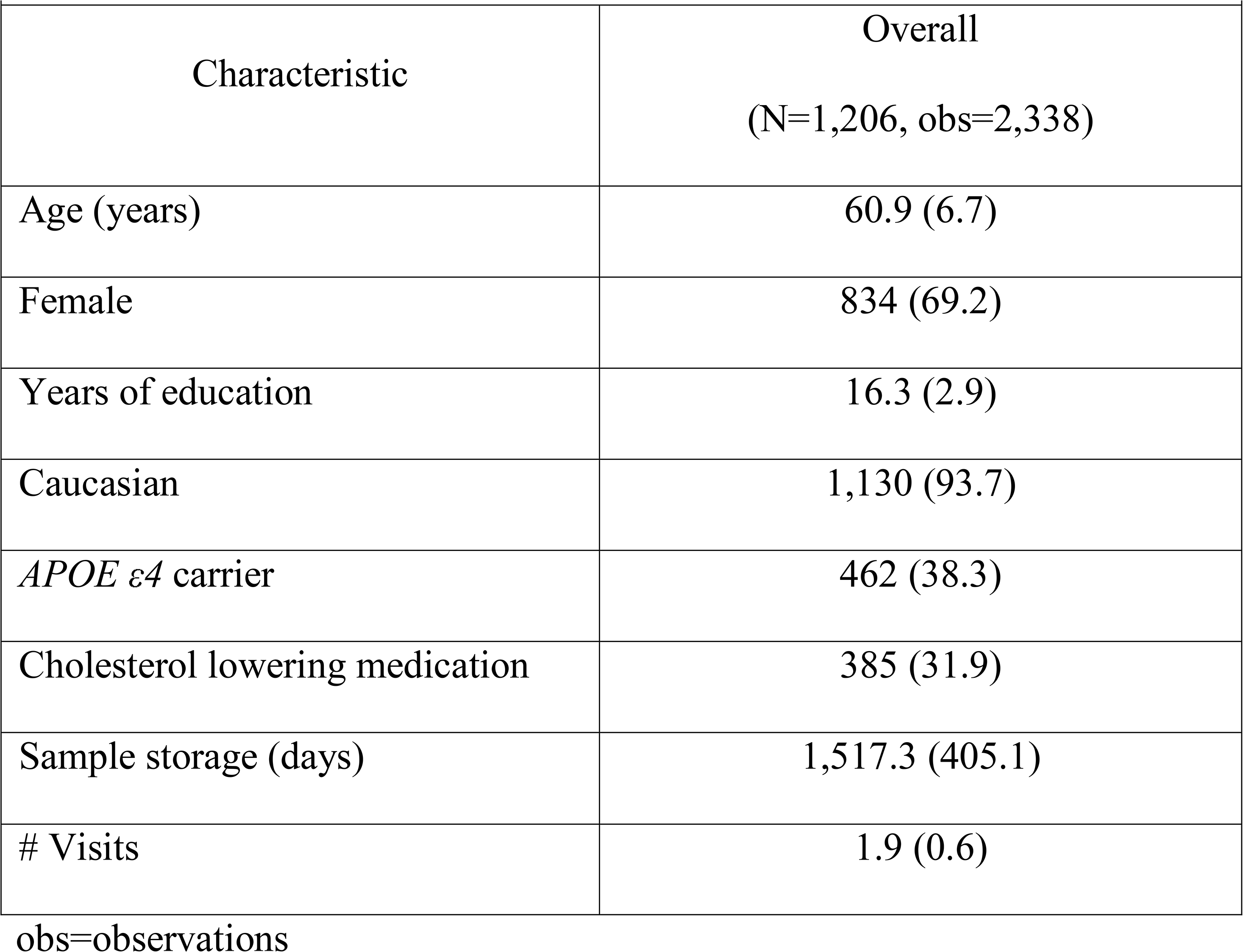
WRAP Participant Characteristics at Baseline. Mean (SD) or N (%).

| Table 1. WRAP Participant Characteristics at Baseline. Mean (SD) or N (%). |  |
| --- | --- |
| Characteristic | Overall<br>(N=1,206, obs=2,338) |
| Age (years) | 60.9 (6.7) |
| Female | 834 (69.2) |
| Years of education | 16.3 (2.9) |
| Caucasian | 1,130 (93.7) |
| *APOE* $\epsilon 4$ carrier | 462 (38.3) |
| Cholesterol lowering medication | 385 (31.9) |
| Sample storage (days) | 1,517.3 (405.1) |
| # Visits | 1.9 (0.6) |

obs=observations

**Table 2.**
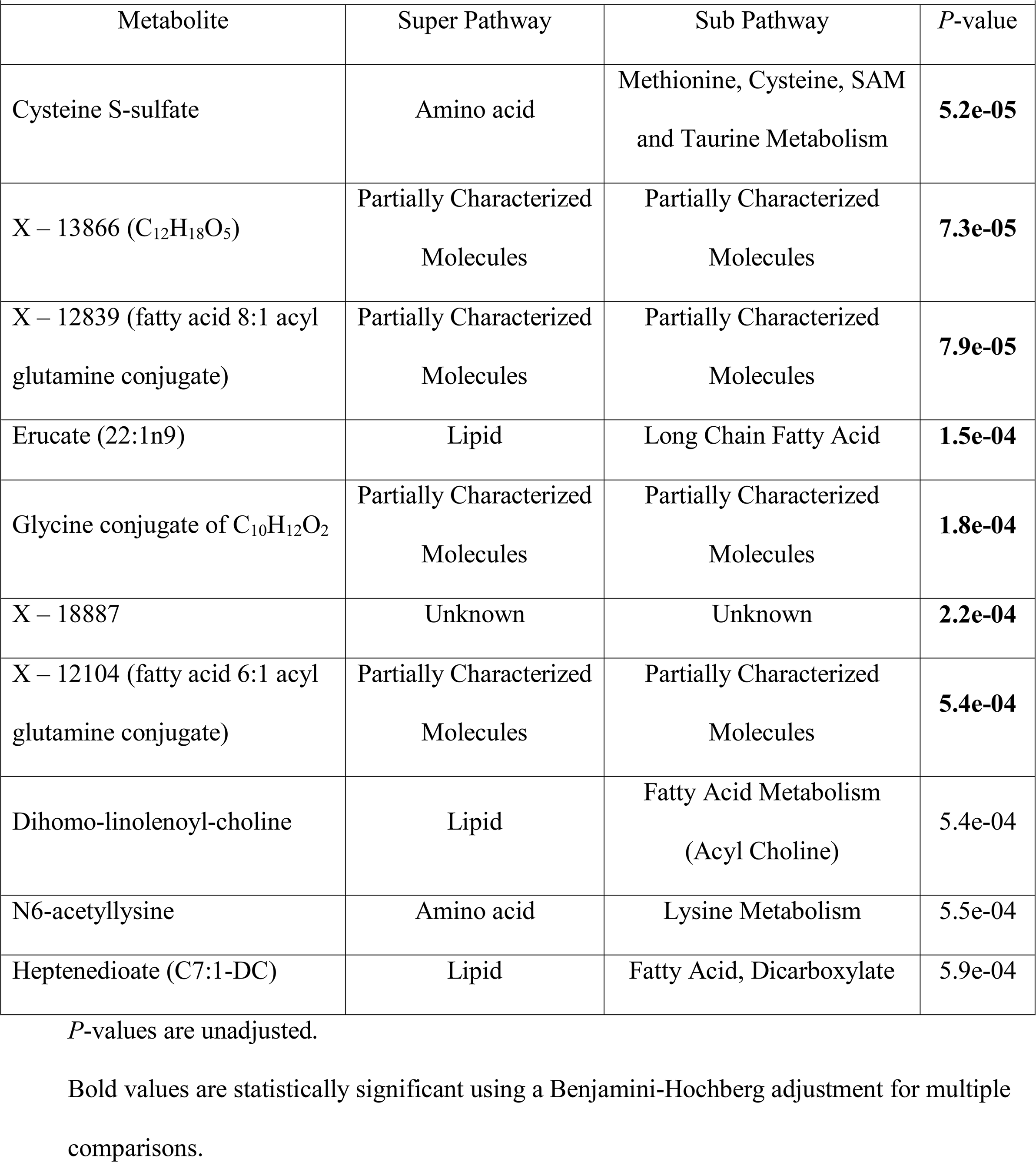
Top ten metabolite*age interactions on executive function

### 3.2. Metabolome-wide association studies

#### 3.2.1. Executive Function

All metabolome-wide association results are detailed in Table S1. Seven metabolite-by age interactions were associated with executive function (Figures 1A and 2). Levels of cysteine S-sulfate, an amino acid, had the strongest association (unadjusted *P*=5.2e-05), with lower levels associated with better executive function in midlife and poorer executive function later in life. The six other significant metabolites showed the opposite relationship with age and executive function, such that lower metabolite levels were associated with poorer executive function in midlife and better executive function later in late. These metabolites included erucate (22:1n9) (a monosaturated omega-9 fatty acid), four partially characterized molecules (glycine conjugate of C_10_H_12_O_2_, fatty acid 8:1 acyl glutamine conjugate, fatty acid 6:1 acyl glutamine conjugate, and C_12_H_18_O_5_), and one unknown metabolite (X-18887).

**Figure 1.**
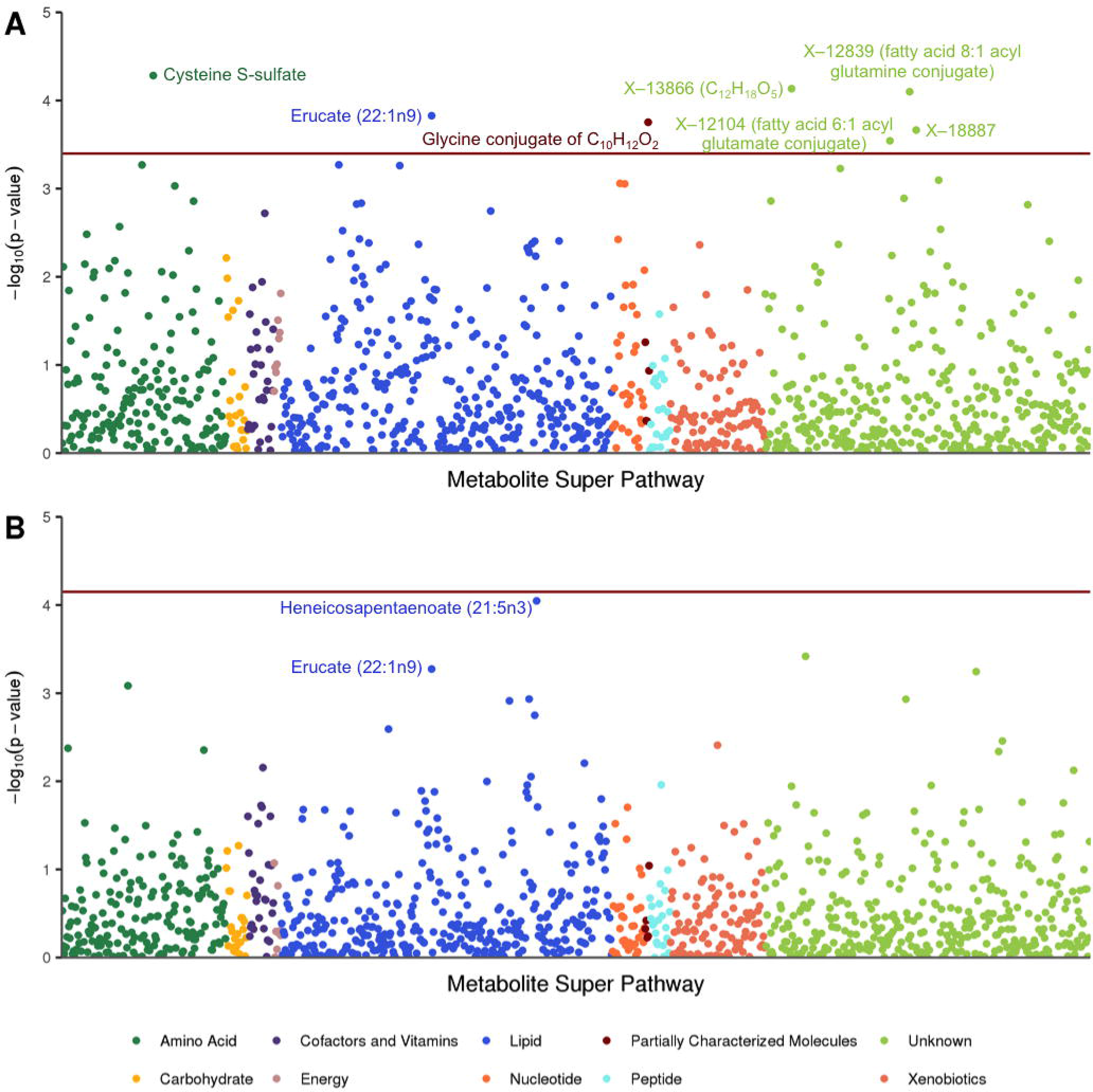
Manhattan plot of metabolome-wide association results for cognitive composite scores. A. Seven metabolite*age interactions were significantly associated with executive function. B. No metabolite*age interactions were significantly associated with delayed recall. Both sets of results used a Benjamini-Hochberg adjusted *P*-value threshold (red horizontal line).

**Figure 2.**
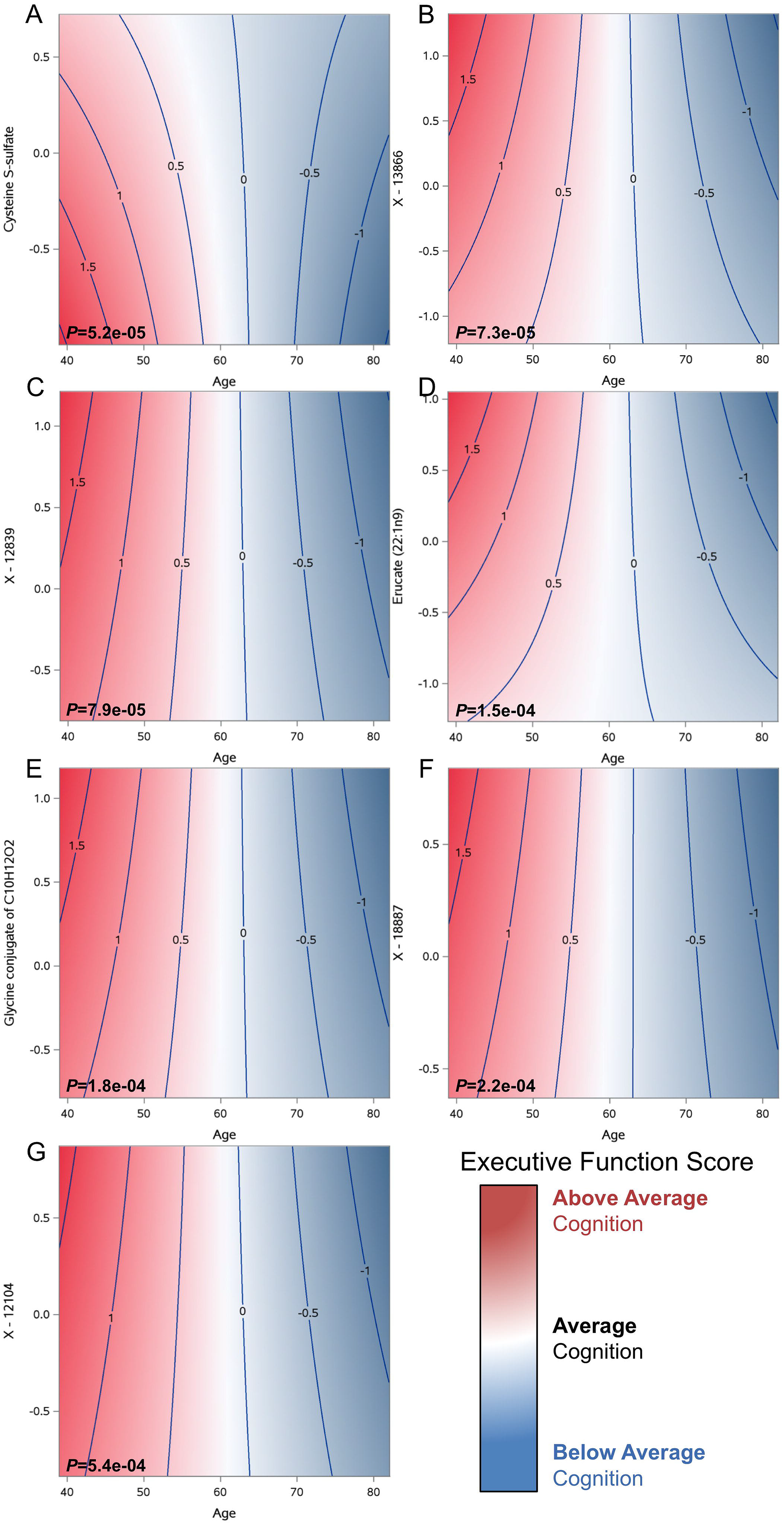
Contour plots showing executive function trajectories by seven time-varying metabolite levels. The x-axis represents age, y-axis represents standardized metabolite levels, and z-axis represents the executive function composite score. In younger ages, higher levels of most metabolites are associated with above average cognition (indicated by the darker red regions), whereas in older ages, higher levels are associated with below average cognition (indicated by the darker blue regions), with the exception of cysteine s-sulfate, where there opposite is true.

#### 3.2.2. Delayed Recall

No metabolite-by-age interactions were associated with delayed recall after adjusting for multiple comparisons. The three strongest interactions included heneicosapentaenoate (21:5n3) (a polysaturated fatty acid, unadjusted *P*=0.00009), X − 02269 (an unknown metabolite, *P*=0.0004), and erucate (22:1n9) (unadjusted *P*=0.0005) (Figure 1B). Four of the seven metabolites associated with executive function showed a similar relationship with delayed recall, although none were statistically significant (erucate (22:1n9), X − 13866, X − 12104, and cysteine S-sulfate, all unadjusted *P*-values <0.20) (Figure S2).

### 3.3. Mendelian randomization

GWAS summary statistics were available for three of the seven metabolites associated with executive function (cysteine S-sulfate, erucate (22:1n9), and X-13866, an unknown metabolite) and used to create a PS for each metabolite. The three PSs were fairly weak instruments, with correlations with corresponding metabolites ranging from r=−0.04 to 0.004 and the largest *F*-statistic=1.71, well below the commonly used *F*-statistic threshold of 10 (Stock et al., 2002) (Table S2). Not surprisingly, associations between executive function and the PSs-by-age were not significant (each *P*≥0.54). Thus, MR analyses were insufficient to draw conclusions about the nature of the relationship between the metabolites and executive function.

### 3.4. Metabolite pathway analysis

Of the 1,097 metabolites tested, only 291 had KEGG compound IDs that were recognized by MetaboAnalyst and could be used as the reference panel for the pathway analysis. A total of 254 metabolites met the inclusion threshold of an unadjusted *P*<0.05 for the cognitive metabolite pathway analysis; however, only 82 of these were identified metabolites with KEGG compound IDs. These metabolites most strongly represented pathways involved in inositol phosphate, ether lipid, and amino sugar and nucleotide metabolism, although none of the pathways identified were statistically significant (Figure S3 and Table S3).

## 4. Discussion

We analyzed the metabolomics of cognitive trajectories using time-varying plasma metabolomic samples and a large panel of metabolites. Our findings suggest that specific metabolite levels, particularly cysteine s-sulfate and fatty acid lipids, correspond with executive function trajectories in late middle-aged adults at increased risk for AD. However, metabolite levels were not statistically associated with delayed recall trajectories.

The associations we observed between metabolite levels and executive function trajectories could provide insight to mechanisms contributing to cognitive decline. In particular, lower levels of the amino acid cysteine S-sulfate were associated with better executive function in midlife, but poorer executive function later in life. The involvement of cysteine metabolism in AD has been implicated in a pathway analysis of previous AD metabolomics studies (Enche Ady et al., 2017). Our results further suggest that such a relationship could depend on age. Cysteine S-sulfate is a glutamate receptor agonist that can lead to calcium influx in nerve cells and neurotoxicity when present in high levels (Olney et al., 1975; Snowden et al., 2017). It has been shown to drive excitotoxic neurodegeneration in individuals with molybdenum cofactor deficiency, an autosomal recessive inborn error of metabolism characterized by early childhood death (Kumar et al., 2017). This supports our finding that high levels of cysteine S-sulfate may be detrimental to cognitive function in midlife. However, further investigations using longitudinal cohorts, and perhaps experiments using model organisms, will be crucial to validate our findings and determine whether high levels could have protective effects later in life.

The opposite pattern was seen for the six other metabolites associated with executive function, which included three fatty acids, where higher metabolite levels in younger years, but lower levels in older years, were associated with better executive function. One of these fatty acids was erucate (22:1n9), an omega-9 fatty acid that readily crosses the blood brain barrier (Golovko and Murphy, 2006) and has been shown to enhance memory performance in mice (Kim et al., 2016). Fatty acids have long been suspected to influence cognitive performance, but studies have had mixed findings regarding their role, particularly of omega-3 fatty acids (Cederholm et al., 2013; Mazereeuw et al., 2012). Our results suggest that this role may be difficult to define because the implications of these metabolite levels change as individuals age. This is further supported by similar relationships we found for two partially characterized fatty acids (fatty acid 8:1 acyl glutamine conjugate and fatty acid 6:1 acyl glutamine conjugate). More information about these two particular metabolites could prove useful in understanding the relationship between fatty acids and cognitive function. Beyond cognitive performance, omega fatty acids have also been shown to be dysregulated in certain brain regions of patients with AD pathology (Snowden et al., 2017), further strengthening the potential relevance of fatty acids. Further, a recent study reported higher levels of docosapentaenoate (22:5 n-6), a long-chain polyunsaturated fatty acid, to be associated with less decline of information-processing speed in a sample of midlife African Americans (Bressler et al., 2017), supporting the association between higher fatty acids levels and better cognition function in midlife.

This study had several limitations. The pathway analysis we performed was highly limited due to the large number of metabolites in our panel that did not have KEGG compound IDs. This greatly underscores the importance of continued efforts to identify and characterize metabolites. The PSs we developed for our MR assessment were weak instrumental variables and did not allow us to determine whether levels of the metabolites we identified are causally related to executive function. Our cohort may not have experienced sufficient cognitive decline at this point to identify metabolite levels associated with delayed recall trajectories. Because our analyses are based on an average of two and up to three longitudinal samples per participant, we are somewhat limited in our ability to assess cognitive and metabolite trajectories. Although our metabolite panel is large relative to previous investigations, it is possible that a different panel of metabolites could produce different results; however, quantifying and identifying metabolites is a challenging task that is highly dependent on technological advances. It will be critical to replicate findings presented here with an independent cohort. To assist with replication of metabolomics findings, as the field of metabolomics rapidly develops, it will be crucial to develop standard methods of measuring and analyzing metabolites.

Using a large panel of longitudinal metabolomics data, we found that levels of certain plasma metabolites, including cysteine S-sulfate and erucate, were associated with age-related change in executive function, one of the earliest aspects of cognitive function to change in the course of AD development. Replication in cohorts with longitudinal metabolomics data will be necessary to confirm whether these metabolites contribute to the development of AD. If these metabolites are shown to have causal influences on cognition through future longitudinal and experimental research studies, subsequent investigations of their nutritional influences could further elucidate the mechanisms influencing early stages of AD and perhaps inform preventative measures.

## Declarations of interests

none

## Acknowledgements

BFD was supported by an NLM training grant to the Computation and Informatics in Biology and Medicine Training Program [grant number NLM 5T15LM007359]. This research was also supported by the NIH [grant numbers R01AG054047, R01AG27161, UL1TR000427, and P2C HD047873], Helen Bader Foundation, Northwestern Mutual Foundation, Extendicare Foundation, and State of Wisconsin. The authors thank the University of Wisconsin Madison Biotechnology Center Gene Expression Center for providing Illumina Infinium genotyping services. GWAS summary statistics for several metabolites excluded from analyses in the Long et al., 2017 Nature Genetics publication were generously calculated and provided by Elizabeth Cirulli, PhD, and authors of the Long et al. manuscript. The authors also thank Joshua Coon, PhD, for providing his expertise to early versions of this investigation. We especially thank the WRAP participants.

